# RNA dicing regulates the expression of an oncogenic JAK1 isoform

**DOI:** 10.1101/2024.02.20.581186

**Authors:** Yuval Malka, Rob van der Kammen, Shinyeong Ju, Ferhat Alkan, Cheolju Lee, William James Faller

## Abstract

mRNA transcripts have limited potential for protein synthesis, defined by their open reading frames^1^. However, recent advances have revealed a far more complex reality, in which the proteome exceeds the perceived limits of the transcriptome^2–6^ exposing a significant gap in our understanding. Our prior studies have demonstrated that mRNA can undergo further processing, yielding truncated, uncapped mRNAs with translation potential^2,7^. Yet, the biological importance of this process remains mostly unclear. Here, we demonstrate that cleavage within the coding sequence of JAK1 mRNA produces an uncapped variant downstream to the cleavage site, at the expense of the full-length transcript. This results in the independent translation of the JH1 kinase domain, a process we term ‘RNA dicing’. Notably, canonical and diced JAK1 variants have distinct impacts on cell proliferation and tumorigenesis, operating independently, localizing to distinct cellular compartments. In addition, the activation of JAK1 signaling through IFNγ induction promotes the dicing of JAK1, thereby altering the balance of isoforms present. Base editor screens reveal that stop-codon nonsense mutations, which are typically considered loss-of-function, have differing impacts depending on their position relative to the dicing site. In agreement with this, JAK1 nonsense frameshift (JAK1fs) mutations in endometrial tumors inhibit the tumor-suppressive functions of canonical JAK1, while amplifying the oncogenic potential of the diced JH1 kinase domain. Lastly, we demonstrate that Momelotinib^8^, a JAK1-specific inhibitor, is more effective in cancer cells carrying a “loss-of-function” JAK1fs mutation, highlighting the significant impact of RNA dicing biology as a potent tool for patient stratification. Our findings characterize RNA dicing as a fundamental regulatory machinery that diversifies the potential products of a single mRNA molecule, and allows for significant variation in biological function.

## INTRODUCTION

Traditionally, genes have been perceived as blueprints for proteins with a distinct function, yet it is clear that individual proteins possess significant variability in their biological roles. These include context-dependent potential for localization, interaction, and other functions^9,10^. Such variability is partially explained by post-translational modifications, or by RNA metabolic processes like RNA splicing, but it remains largely undefined for most proteins. Proteomic analyses, enhanced by high-resolution mass spectrometry, has shown that the diversity of proteins expressed in cells significantly exceeds predictions based on gene count^2–4^. This inconsistency highlights a greater complexity in our proteome than expected, and suggests gaps in our knowledge of protein synthesis^11,12^. Indeed, recent advances in this field have revealed extensive translation events beyond canonical open reading frames^1,13^ which may help resolve this apparent disagreement.

Previously, we reported a widespread phenomenon whereby mRNAs from thousands of genes undergo endonuclease cleavage, producing truncated 5’ Uncapped Polyadenylated Transcripts (5’UPT) with translation potential. This production of multiple functional isoforms from a single mRNA can significantly diversify the proteome^2,7^, potentially resolving some of the discordance between gene number and protein number.

Here, we take this further, and show that such mRNA cleavage functions to produce transcripts encoding specific subsets of protein domains, and the resulting proteins have distinct biological functions. We demonstrate that cleavage of the JAK1 mRNA produces a transcript encoding just the C-terminal kinase domain. This results in a protein that lacks both the FERM domain (which is crucial for membrane localization), and the pseudokinase JH2 domain (which is known to negatively regulate the kinase domain). The resulting novel protein isoform shows altered localization, as well as enhanced kinase activation compared to the canonical protein. This process, which we term ‘RNA dicing’, represents a new model by which the metabolism of mRNA allows the production of multiple proteins from one gene, highlighting that an mRNA molecule is not a monolithic biological subunit, but the template for discrete, functionally distinct, products.

## RESULTS

### Dynamic Expression of Protein Domains Through RNA Dicing

We have previously shown that endonuclease cleavage of RNA at an alternative polyadenylation site can generate uncapped, autonomous RNA fragments with translational potential^2,7^. This process requires several features, including APA-mediated cleavage, structured RNA to stabilize the new 5’-end, and m6A modifications to facilitate cap-independent translation. Notably, increased APA usage, which is prevalent in various biological contexts, amplifies RNA dicing, and is predicted to favor the production of 5’ UPTs over canonical mRNA variants. The truncation of an mRNA within the open reading frame^1^ would therefore result in a translated protein containing only the specific domains contained in the new, truncated mRNA isoform. We hypothesized that RNA dicing could selectively exclude certain protein domains, creating new isoforms with potentially novel biological properties (Fig.1a). Indeed, it is well known through domain-specific deletion assays carried out across numerous studies and genes that protein domains function differently depending on their context, via altered localization or the absence of inhibitory domains, for example^14–16^.

We therefore analyzed multi-omics datasets (including APA sites, uncapped 5’end RNAseq, m6A RIPseq, ribosome profiling following harringtonine treatment, and mass spectrometry) to identify translation initiation sites, and potential diced mRNAs that may display altered function. This analysis revealed many examples of such mRNAs. For example, the LUC7L3 mRNA is predicted to be cleaved, resulting in the independent expression of the arginine-serine-rich sub-domain, which has previously shown to maintain interaction capability when artificially independently expressed^14^ (Extended Data Fig.1a,b). Next, we focused our search on the kinase protein family. Multiple studies have shown that kinase catalytic domains can exhibit functionality when independently expressed^15,17,18^. Therefore, we globally assessed whether RNA dicing preferentially produces independent kinase domain expression. Combining m6A-RIPseq data and N-terminal Mass Spectrometry (Nterm MS) to identify internal non-canonical translation start sites potentially produced by dicing, we observed a clear m6A peak directly preceding kinase domains, which was absent in non-kinase domains of the same genes. This suggests that RNA dicing may not occur at random sites and may induce functional independent kinase expression post-RNA dicing (Fig.1b).

In order to understand the biological consequences of this apparent selective production of independent kinase domains, we focused on the JAK1 kinase, a member of the JAK kinase family. These intracellular, non-receptor tyrosine kinases are pivotal in signaling transduction following exposure to various cytokines and growth factors^19^. The JAK family consists of four members, each characterized by four major domains^20^. The JH1 domain is the active kinase domain responsible for enzymatic activity. In contrast, the JH2 domain functions as a pseudokinase, regulating the activity of the JH1 domain^15,18^. The other two domains, FERM and SH2, are crucial for binding to cytokine receptors and regulating kinase activity (Fig.1c).

Multi-omic analysis of JAK1 in HeLa cells revealed a conserved polyadenylation signal in exon 8, aligning with the C-terminal of the FERM domain. Active APA usage at this site was confirmed using 3’ polyadenylation sequencing (Fig.1c). Both digestion of uncapped mRNA by Terminator TM 5’-Phosphate-Dependent Exonuclease (TEX)^5^ treatment, and 5’ uncapped RNAseq, confirmed accumulation of uncapped variants downstream to the APA cleavage site (Fig.1c, Extended Data Fig.2b). As has been previously described, uncapped mRNAs that are highly structured at their 5’end can avoid exonuclease degradation^2,21^. We therefore analyzed icSHAPE data^22^, which indicated a highly structured 5’end downstream to the APA site, corresponded to the 5’end of the 5’UPT. In addition, cytoplasmic icSHAPE analysis showed higher structural integrity than nuclear fractions, suggesting post-transcriptional cleavage (Extended Data Fig.2a). Long-read Nanopore sequencing further supported the existence of JAK1 5ʹUPTs, and these also decreased following TEX treatment (Fig.1d). m6A RIPseq identified modification sites within the JAK1 CDS, and Nterm MS analysis indicated two non-canonical translation sites, one starting directly downstream of the APA site and a second corresponding to the strongest m6A site signal (Fig.1c). GROseq^23^ and H3K4Me3^24^ analysis ruled out the possibility that these 5’UPTs derive from non-canonical transcription (Extended Data Fig.2c). Interestingly, transcript annotations of JAK1 predict a C-terminal variant corresponding to the JH1 kinase domain. This finding, which can be explained by RNA cleavage (Fig.1c lower panel), further supports our observations.

To assess the production of the diced JAK1 mRNA at the protein level, we performed western blot analyses using antibodies targeting either the JH1 kinase domain or an upstream region (aa 551-776) corresponding to the JH2 domain. The JH2-specific antibody revealed the expected full-length JAK1 protein at 120 kDa (Fig.1e). However, the JH1 antibody detected additional bands around 37 kDa (Fig.1e). Given that JAK1 phosphorylation occurs at the JH1 domain, we further probed cell extracts with two different antibodies, both raised against phosphorylated JAK1. These identified the same two bands around 37/39 kDa (Fig.1e). Lastly, we used siRNA to knock down either the JH1 or JH2 domains of JAK1, and measured the effect using primers targeting up-stream of the cleavage site (full length JAK1), or down-stream of the cleavage site (full length JAK1 and diced JAK1). In the presence of the diced isoform, we would expect the siRNA targeting the JH1 domain to have a larger effect on the downstream product (full-length JAK1 and diced JAK1) compared to the upstream product (full-length JAK1). In line with the previous results, this is precisely what we observe: the siRNA targeting the JH1 domain has a significantly greater effect on the downstream product compared to the upstream (Fig.1f). At protein level, treatment with the JH2 and JH1 siRNAs resulted in the anticipated decrease of canonical JAK1. Surprisingly, siRNA targeting JH2 increased the expression of the diced product, potentially indicating an induction of additional dicing following canonical JAK1 knock down. siRNAs targeting JH1, on the other hand, had the expected effect on the canonical isoform but a minor effect on the JH1 domain (Extended Data Fig.2d). Our findings indicate that RNA dicing of JAK1 leads to the production of truncated mRNA, potentially producing an independent JH1 kinase domain.

### RNA Dicing plays a critical role in Determining JAK1 Isoform Function

To study the potential biological consequences of independent JH1 domain expression we designed three constructs, and expressed them in the MCF7 cell line: 1) wild-type (WT) full length JAK1; 2) APA-mutant JAK1 (in which we mutated the poly-adenylation sequence without changing the amino acid sequence), and 3) codon-optimized JAK1 (in which we mutated 20% of the nucleotide sequence across the entire JAK1 CDS, while maintaining the WT amino acid sequence) (Fig.2a). To assess the efficiency of dicing across different constructs, we designed two pairs of construct-specific qRT-PCR primers that do not detect endogenous JAK1: the first targeting upstream of the dicing site; and the second downstream of the dicing site. We then used TEX treatment to digest the diced, uncapped RNA, and measured the abundance of the upstream and downstream products, normalized to the relevant untreated control. As expected, the WT construct showed substantially decreased downstream product, indicating that a significant portion of that product derives from the TEX-sensitive diced isoform. However, the downstream products in both the APA-mutant and codon-optimized constructs show decreased sensitivity to TEX, indicative of reduced JAK1 mRNA dicing, and consequent reduced expression of the diced isoform of JAK1 (Fig.2b).

At the protein level, western blot analyses indicated increased full-length JAK1 expression and reduced phosphorylation of the JH1 kinase domain presence in these constructs, supporting a role for dicing in JAK1 phosphorylation (Fig.2c). Interestingly, both APA-mutant and Codon-optimized constructs showed higher levels of non-phosphorylated, truncated JAK1 which may derive from increased endogenous expression of JAK1 (Fig.2c). qRT-PCR targeting regions upstream and downstream of the dicing site (FERM and JH1 domains respectively), specific to endogenous JAK1 mRNA, revealed that the codon-optimized construct increased endogenous JAK1 mRNA levels, particularly evident using primers specific to the JH1 domain, suggesting increased dicing of the endogenous JAK1 following expression of the dicing-resistant construct (Fig.2d). To further address the dynamics of canonical and diced JAK1 expression, we treated the cells with IFNγ, which activates the JAK-STAT pathway via JAK phosphorylation. Interestingly, we observed higher phosphorylation levels in both the canonical and diced JAK1 isoforms in the control non-transfected and WT JAK1 contexts compared to APA-mutant and Codon-optimized contexts (Fig.2c). These results indicate that a change in dicing may interfere with the JAK1 signaling cascade, and the phosphorylation of both canonical and diced JAK1.

To explore this possibility, we investigated the dicing of JAK1 mRNA in macrophage activation^25,26^, given the well-recognized crucial role of JAK-STAT signaling in IFNγ-mediated activation of these cells. Upon differentiation of HL-60 cells to M1 macrophage (using IFNγ and LPS), there was a shift from predominantly intact non-diced JAK1 mRNA variants (enabling canonical JAK1 signaling) to diced variants (Fig.2e). Western blot analysis validated the change in the expression balance between canonical and diced JAK1 following HL-60 differentiation to M1 macrophages (Fig.2f). These results suggest that mRNA dicing of JAK1 is dynamic and depends on the biological context. Moreover, the robust effect on JAK1 dicing and the independent expression and phosphorylation of JH1 following IFNγ treatment (a well-known inducer of JAK1 signaling) strongly indicates that this process plays a significant role in the JAK1 signaling cascade.

Lastly, we assessed the biological impact of the inhibition of JAK1 dicing. JAK kinases are known for their dynamic cellular localization, and opposing effects in different types of cancer^27^. JAK1 is found in both the cell membrane and nucleus, with its nuclear presence linked to rapid cell cycle progression and tumorigenesis^28,29^. In addition, nuclear JAK1 has been shown to regulate gene expression not only by activating transcription factors other than the STATs, but also through epigenetic activities, such as the phosphorylation of histone H3^30^. Interestingly, early studies reported that phospho-JAK1 (which appears as short isoform) was present in the nuclear fraction while full length JAK1 was present in the cytoplasmic fraction^31^. Western blot analysis of cytoplasmic and nuclear fractions validates this previous observation, indicating that diced JH1 and canonical JAK1 localize to different cellular compartments (Extended Data Fig.2e).

Given that previous studies have indicated nuclear JAK1 expression leads to increased cell proliferation, we examined the effect of JAK1 dicing on proliferation, using our JAK1 constructs. Cell proliferation assays in MCF7 cells showed that APA-mutant and Codon-optimized had significantly reduced cell proliferation (Fig.2g), a finding that was mirrored in colony formation assays (Fig.2h, Extended Data Fig.2f). These findings underscore a critical interplay between canonical full-length JAK1 and truncated JH1 domain expression, mediated by JAK1 dicing, which in turn influences cellular proliferation rates.

### Impact of Genomic Alterations in JAK1 on Cancer Progression Through RNA Dicing

Genetic mutations play a critical role in cancer, with the location of these mutations often dictating whether a gene exhibits a loss of function (LOF) or gain-of-function (GOF). JAK1 mutations are found in 1.88% of all cancers, and are particularly prevalent in specific cancer types such as endometrial, breast, prostate, colon, and lung adenocarcinomas^32^. Notably, JAK1 frameshift (JAK1fs) is thought to be a LOF mutation, and is frequently observed in endometrial cancer^33^ (EC; 10% in TCGA cohort, 13% in CPTAC cohort), with a strong association with microsatellite instability (MSI)^33,34^. Interestingly, while the presence of MSI often predicts a good outcome, JAK1fs has been shown to correlate with higher tumor grades in these cases (Fig.3a). In addition, proteomics analysis of tumors with JAK1fs mutations shows an increase in cell cycle markers and proliferation (Fig.3b).

Mutation mapping indicates that the majority of JAK1fs mutations occur at position 860, situated between the JH1 and JH2 protein domains. Typically, a frameshift mutation such as this would be viewed as a LOF mutation (Fig.3c). However, when RNA dicing is taken into account, it raises the possibility that the JAK1fs mutation might have a different effect on each isoform, silencing the expression of the canonical isoform, while not affecting the diced isoform. The result would be a significant shift in isoform expression, potentially increasing proliferation and other oncogenic processes, as shown in Fig. 2. In effect, this suggests that what is typically considered a LOF mutation may in fact be a GOF mutation, through the selectively silencing one isoform of JAK1.

**Figure 1.**
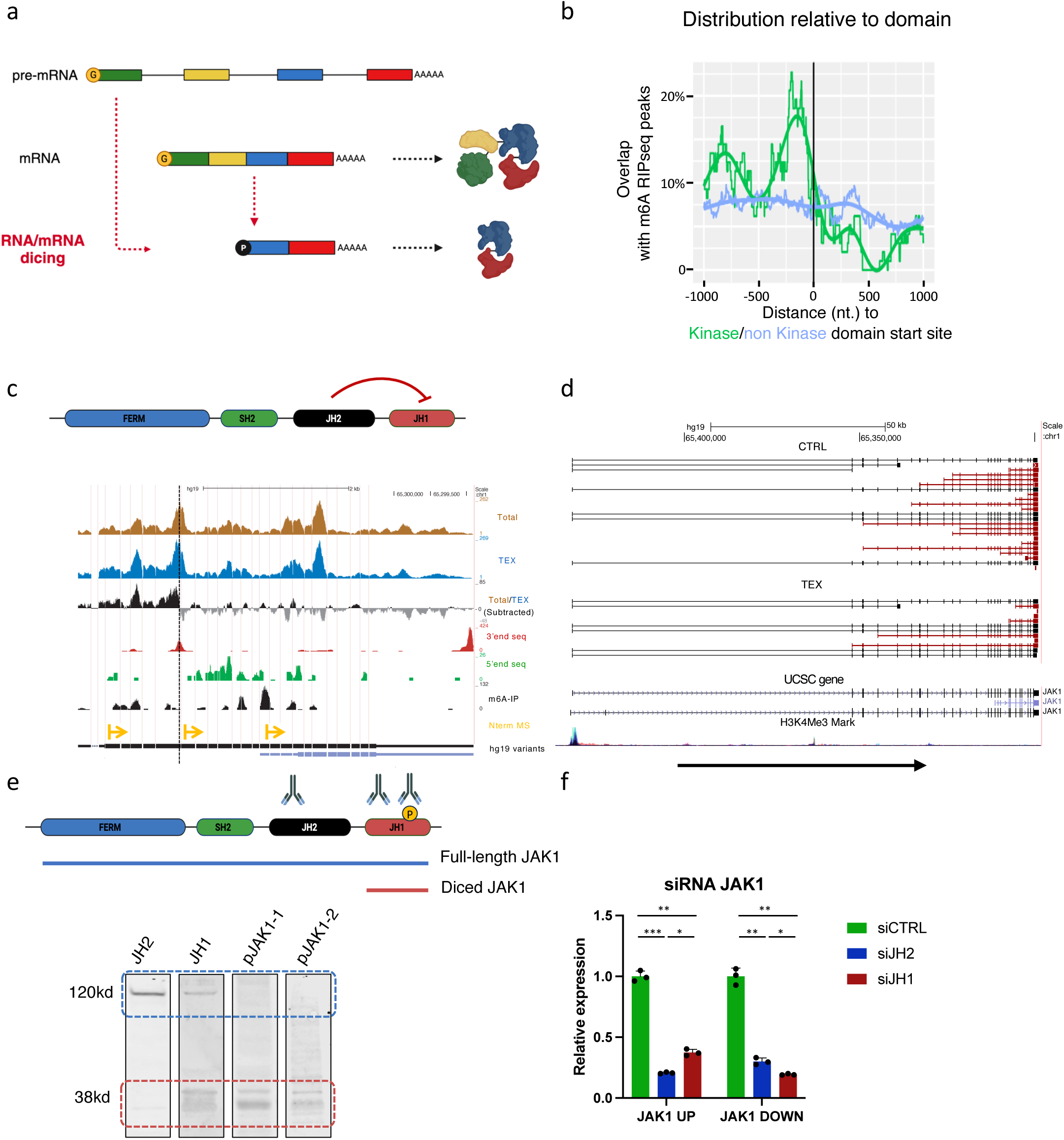
RNA Dicing: Metabolic Mechanism for RNA Processing. (a) **Schematic** model illustrating how endonuclease cleavage triggers the expression of downstream transcript isoforms. These isoforms, stabilized by RNA structure, serve as templates for uncapped non-canonical protein synthesis. (b) m6A modifications are enriched near kinase domain boundaries. Overlap of m6A-RIPseq peaks with kinase and non-kinase domain boundaries. The x-axis represents the distance from reference positions on the mRNA level, while the y-axis shows the proportion of m6A-RIPseq peaks overlapping at those positions. Lines of various colors in the figure correspond to different reference positions, categorizing them as either kinase or non-kinase domain start positions. (c) Schematic representation of the JAK1 domains: FERM - a protein module involved in localizing proteins to the plasma membrane; SH2 - commonly found in adaptor proteins aiding in the signal transduction of receptor tyrosine kinase pathways; JH2 (Pseudokinase) - a catalytically-deficient pseudoenzyme; JH1 (Kinase) – a module with catalytic function in protein kinases. JH2 induces inhibition. RNA-seq read coverage (y-axis) across the putative cleavage point (dash line) of JAK1 shows reduced downstream read coverage in TEX (exonuclease) compared to coverage in untreated RNA (Total). The Total/TEX subtracted panel (black) exhibits a sharp decrease in RNA levels between exons 8/9. APA (3’end seq), 5’end seq, and m6A-IP show endonuclease cleavage at exon 7, resulting in termination/generation of uncapped truncated RNA with translation potential mediated by m6A. N terminal MS shows non-canonical peptides correspond to the cleavage site and to the highest peak of m6A modification site. (d) Nanopore long RNA sequence data genome browser view of isoforms that aligned to JAK1. Black isoforms adjacent to the promoter region at their 5’end, and red isoforms are 5’end diced RNA. Red isoforms show high sensitivity to TEX treatment, indicating they are uncapped, while black isoforms were not affected by the treatment. (e) Western blot analysis for JAK1 using antibodies against JH2 domain, JH1 domain, and two different phospho-JAK1 (JH1) shows expression of different JAK1 isoforms corresponding to full length (blue) or diced JAK1 (red) isoforms. Phospho-JAK1 shows detection of only diced JAK1 kinase domain isoform. (f) Knockdown experiment against JAK1 using two different siRNA (targeting JH1 and JH2 domains) shows a different effect on JAK1 RNA levels at different positions (UP amplicon; FERM domain, DOWN amplicon; JH1 domain). \**p* < 0.05, \*\**p* < 0.01 and \*\*\**p* < 0.001 (two-tailed Student’s *t*-test).

**Figure 2.**
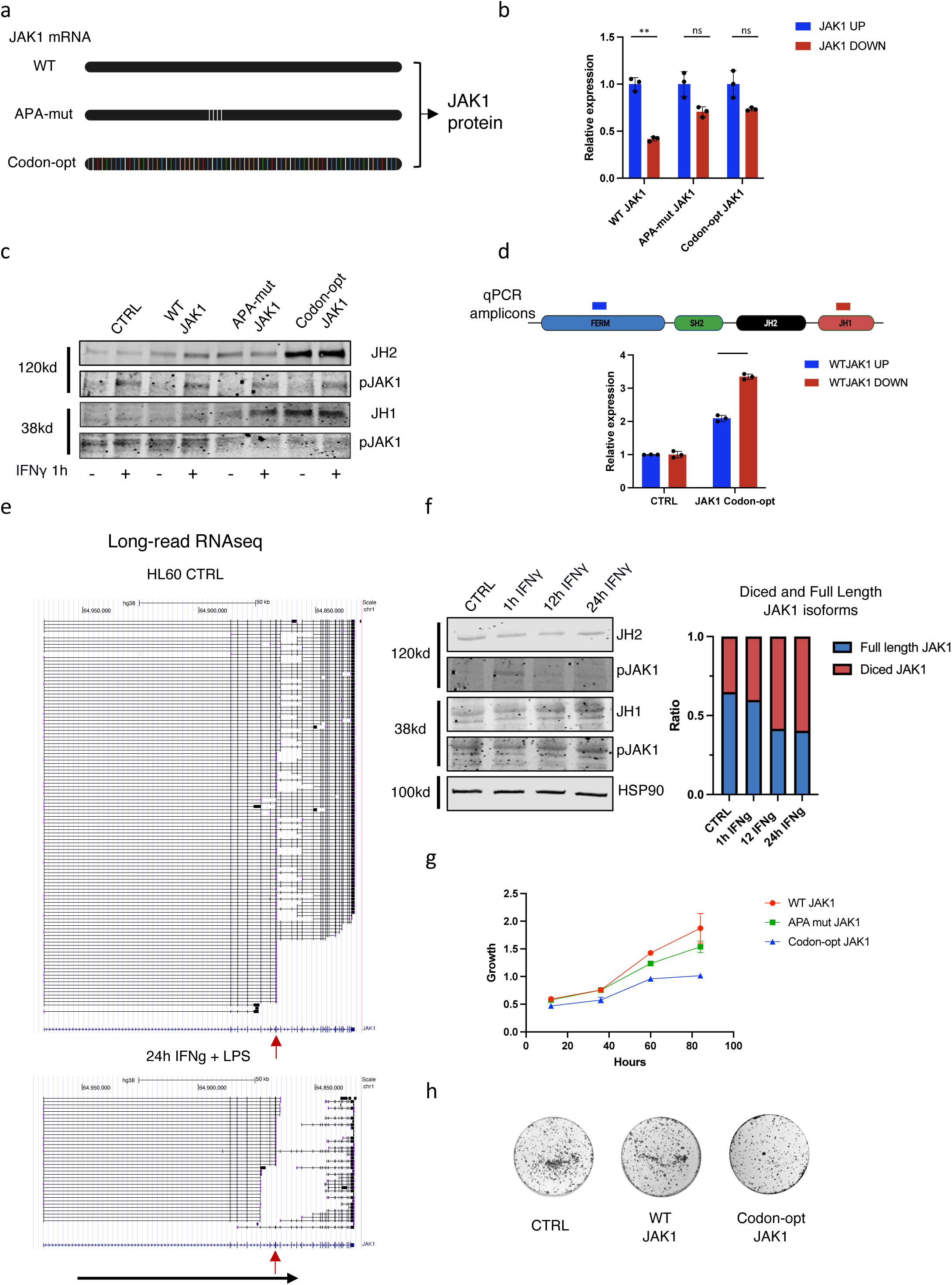
RNA Dicing plays critical role in JAK1 kinase function and cell proliferation. (a) Design of three constructs of JAK1: WT JAK1 - human JAK1 CDS; APA-mut JAK1 - codon optimization of nucleotides 990-1176 in the CDS corresponding to the APA region; Codon-optimized JAK1 - substitutions of ∼20% of CDS nucleotides. All constructs contain the open reading frame of WT JAK1. (b) qRT-PCR analysis was performed on the upstream and downstream regions of the JAK1 cleavage site following TEX treatment. The relative expression is normalized to RNA levels prior to treatment. \*\**p* < 0.01 (two-tailed Student’s *t*-test). (c) Western blot analysis of different expression of JAK1 constructs. Codon optimization of JAK1 interferes with JAK1 mRNA dicing, resulting in differential expression of full-length canonical JAK1 and diced phospho-JAK1. Detection was conducted on the same samples using an intact blot with different secondary antibodies or partial stripping to assess the internal JAK1 isoform expression ratio. (d) RT-qPCR of Codon-optimized JAK1-expressing cells shows high expression of endogenous WT JAK1 mainly in diced JAK1 mRNA. Data is normalize to house keeping gene. \*\**p* < 0.01 (two-tailed Student’s *t*-test). (e) Long-read RNAseq of JAK1 in HL-60 cells before and after applying 100ng/ml IFNγ and LPS treatment for 24h. Red arrow indicates the cleavage site. (f) Western blot analysis of JAK1 was conducted using different antibodies at various time points following IFNγ and LPS treatment. On the right, the relative ratio of canonical JAK1 (120 kDa) to the diced, non-phosphorylated isoform (JH1 domain) is presented for these time points. (g) Cell growth of cells expressing three different JAK1 constructs was measured using quantification of crystal violet staining. JAK1 RNA dicing inhibition significantly reduced cell growth and is correlated with high full-length JAK1 protein expression and negatively correlates with diced phospho-JAK1 expression. (h) Colony formation assay shows reduction in colonies in codon-optimized JAK1-expressing cells.

To explore this further, we analyzed three single base editing CRISPR-Cas9 screens designed to introduce stop codons in JAK1 in the HT-29 cell line^35^ (BE3-NGG; BE3.9max-NGN and BE4max-YE1-NGN). Overall, the libraries contained 86 gRNA which introduce stop codons across JAK1 CDS (Table 1), and this data is coupled with a proliferation score. Analysis of this data revealed that stop codons induced outside the kinase domain resulted in increased cellular proliferation, while those within the domain did not affect proliferation rate (Fig.3d, Extended Data Fig.3a), supporting the idea that the kinase domain has a function independent of the full-length protein. Inducing stop codons just downstream of the JH1 domain led to a significant increase in proliferation, possibly due to diminishing inhibitory domain motif known for protein-protein interaction with SOCS1 which regulates JAK1 function. These findings offer additional support for the hypothesis that differential JAK1 expression, driven by RNA dicing, regulates distinct biological functions attributed to different isoforms.

### JAK1fs Mutation Alters JAK1 Function and Increases Malignancy in Endometrial Cancer

It is standard in transcriptome and proteome analysis to normalize reads/peptides to known gene isoforms. As RNA dicing produces new isoforms that overlap with the canonical variant, it is challenging to identify this using standard analysis. We therefore used TCGA data to assess the expression of each domain of JAK1 in endometrial carcinoma patients carrying JAK1860fs mutation. While patients with WT JAK1 exhibited similar expression across all JAK1 domains, JAK1860fs EC patients displayed significant fluctuation in RNA expression for each domain. These samples showed a marked decrease in RNA coverage at the JH2 domain, and a sharp increase in coverage at the JH1 kinase domain. This dip in read coverage strongly suggests discontinuous RNA molecules between the SH2 and JH1 domains, indicating independent expression of the JH1 domain in these patients (Fig.4a). Consistent with this finding, when we then analyzed the protein levels of each JAK1 domain in EC patients using the CPTAC cohort, we observed a similar trend, with a decrease in JH2 MS peptide coverage compared to the SH2 domain and an increased coverage in the JH1 domain indicating translation in frame downstream of the JAK1fs site (Fig.4b).

To account for the potential heterogeneity in tumor JAK1 genomic status, we validated our findings in cell lines with homozygous JAK1860fs mutations, which represent complete depletion of canonical JAK1 expression. High coverage MS analysis of breast carcinoma CAL51 cell line^36^ (JAK1860fs homozygote) revealed over 50% coverage of peptides in the JH1 domain downstream to the homozygous frameshift mutation, confirming in-frame translation past the frameshift site (Fig.4c, Table 2). Western blot analysis of the ISHIKAWA EC cell line (JAK1860fs homozygote, Extended Data Fig. 3b) did not detect full-length JAK1 in either phosphorylated or non-phosphorylated states (Fig.4d). However, the 37kd phospho-JAK1 band was clearly present in both the ISHIKAWA and control MCF7 cell lines (Fig.4d). MS analysis revealed in-frame peptides downstream of position 860^36^, confirming in-frame translation at the JH1 domain in the ISHIKAWA cell line (Table 2).

To examine the potential oncogenic effect of altered JAK1 isoform expression, we restored WT JAK1 expression in ISHIKAWA cell line using constructs expressing either JAK1 WT or JAK1 Codon-opt (Extended Data Fig.3c). Expression of both WT and Codon-opt constructs resulted in decreased growth (Fig.4e). Furthermore, a colony formation assay corroborated these findings, showing stronger decrease in colony formation in Codon-opt expressing cells (Fig.4f, Extended Data Fig.3d). These results support the hypothesis that canonical JAK1 plays a tumor suppressive role, while diced JAK1 may act as an oncogene.

Lastly, we performed a single-sample gene set enrichment analysis comparing mutant JAK1 to WT JAK1 in EC cell lines (23 wild type vs. 10 mutated JAK1). Notably, the most significant decrease in gene set clusters was observed in the negative regulation of macrophage activation, a process that is strongly correlated with JAK1 activation, suggesting higher JAK1 activity in mutant cells, rather than lower (Extended Data Fig.4a). In addition, we corroborated this finding in JAK1fs patient samples (CPTAC cohort), which showed a gene expression signature consistent with M1 differentiated macrophages compare (Extended Data Fig.4b). The increased expression of the diced JH1 kinase domain compared to the canonical JAK1 following nonsense mutations underscores the biological importance of JAK1 dicing, and further supports our hypothesis that JAK1 dicing and JH1 isoform expression play a critical role in cell cycle progression and macrophage activation as shown in Fig 2.

### JAK1 Inhibitor shows Enhanced Efficacy in Cells with JAK1fs Mutations

As the activation of JAK1-mediated signaling plays a key role in tumor invasion and metastasis, there has been a major effort to design therapeutic interventions^37^. While the FDA has approved MSI as a marker of immunotherapy for solid tumors, JAK1fs patients have been shown to be deficient in the antigen processing and presentation, and are likely to be resistant to treatment with immune checkpoint inhibitors^33^. Interestingly, all currently approved small molecule JAK1 inhibitors target the kinase JH1 domain^38^. A previous study has shown that the administration of a JAK1 inhibitor significantly reduces the catalytic activity of the independent JH1 domain compared to the canonical isoform^15^. As the diced state of the JAK1 mRNA may affect the efficacy of JAK1 inhibitors, we next assessed the sensitivity of various cells to JAK1 inhibition. In particular, we focused on CYT387 (Momelotinib), an FDA-approved drug used for the treatment of myelofibrosis^39^, which targets the JH1 kinase domain (ATP competitor with IC_50_ of 11nM).

First, we correlated the efficacy of CYT387 with JAK1 expression in various cell lines. Omics analysis across a range of concentrations (600pM to 38nM) showed no correlation with JAK1 mRNA levels (434 cell lines) or JAK1 protein levels^35^ (221 cell lines; Extended Data Fig.5a, b), emphasizing the poor prediction of pharmaceutical potential using a conventional approach. Next, we selected all endometrial carcinoma cell lines expressing wild type JAK1 or homozygote frameshift mutation (9 and 5 cell lines respectively, Cancer Cell Line Encyclopedia (CCLE)). Strikingly, JAK1fs cell lines are significantly more sensitive to CYT387 compared to the WT expressing lines, at a concentration significantly lower than the published IC_50_ (2.4nM) (Fig.5a). To rule out potential off-target effects and redundancy resulting from other JAK family kinases (JAK2, TYK2, and JAK3, with IC50 values of 18 nM, 17 nM, and 155 nM respectively), we observed no difference in the levels of these proteins between WT and JAK1fs cells (JAK3 protein was not detected, Extended Data Fig.5c). To assess the specificity of the treatment to JAK1 inhibition, we preformed single-sample gene set enrichment analysis of JAK-STAT pathway following CYT387 treatment (GO: Negative regulation of receptor signaling pathway via JAK-STAT). EC cell lines carrying the JAK1fs mutation exhibited a positive correlation between sensitivity to CYT387 (2.4 nM) and negative regulation of the JAK-STAT pathway, a correlation not observed in cells expressing WT JAK1 at this concentration. This further validates the specific inhibitory effect of the JAK1 inhibitor CYT387 on the JAK1 diced kinase isoform (Fig. 5b). Furthermore, JAK1fs is more predictive of sensitivity to CYT387 than almost any other damaging mutation in these cell lines (Fig.5c). All other damaging mutations that have shown higher drug sensitivity in different cell lines have been found to be associated with additional JAK1 damaging mutations.

**Figure 3.**
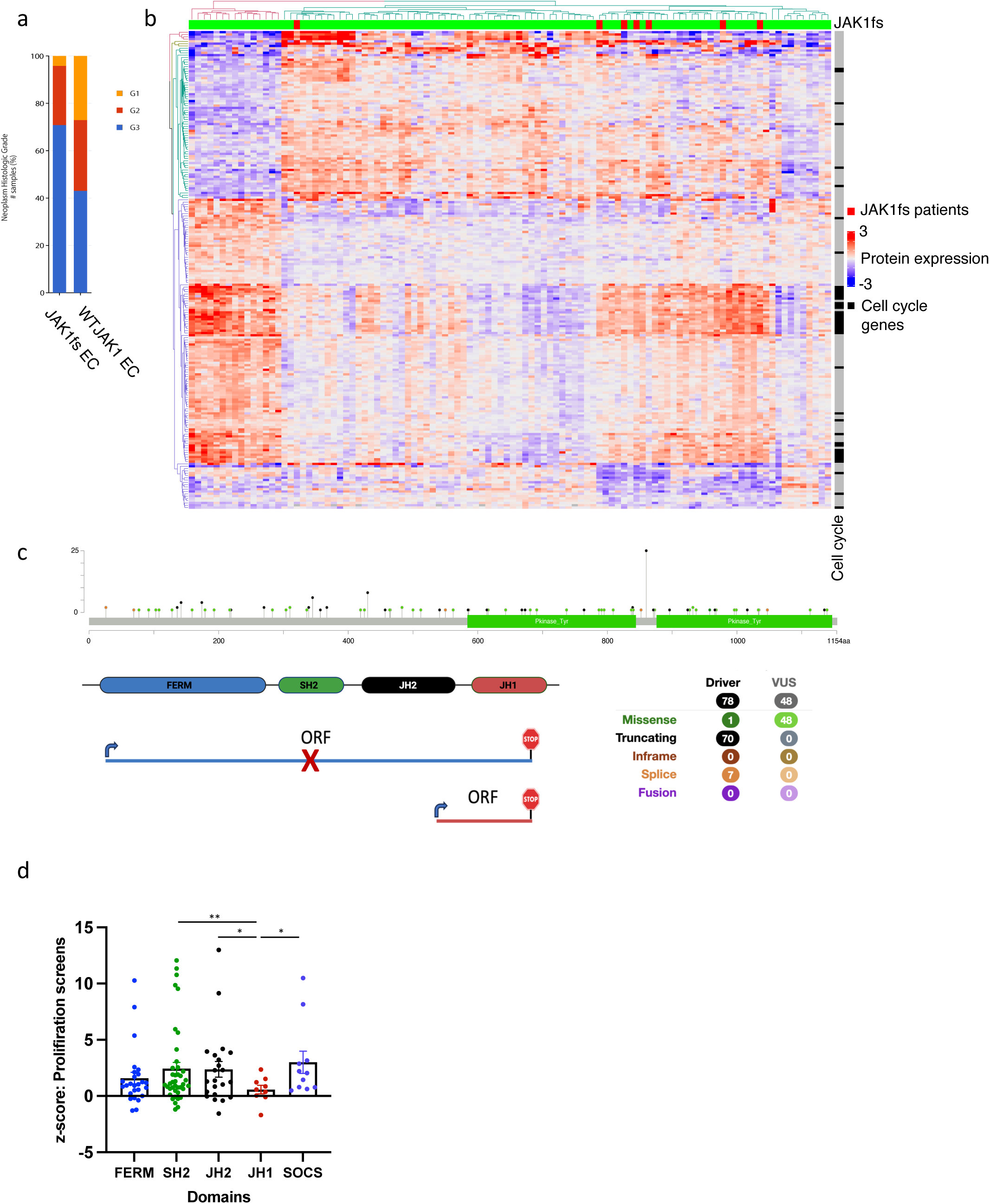
Nonsense/frameshift mutation have different effect on JAK1 expression due to modular expression of canonical and diced isoforms. (a) Tumor grade of Endometrial Carcinoma (EC) in patients with JAK1 frameshift mutations or WT JAK1 expression (CPTAC cohort). (b) Proteogenomic cohort of 95 EC patients (CPTAC cohort) focusing on cell cycle genes. (c) JAK1 hotspot frameshift mutations occur at position 860 (K860Nfs), which are located between the JH2 and JH1 coding regions. (d) Nonsense mutations are separated for each corresponding protein domain mRNA region. Induction of a premature stop codon results in an increased proliferation rate, except for insertions of stop codons at the kinase domain. \**p* < 0.05, \*\**p* < 0.01 (two-tailed Student’s *t*-test).

**Figure 4.**
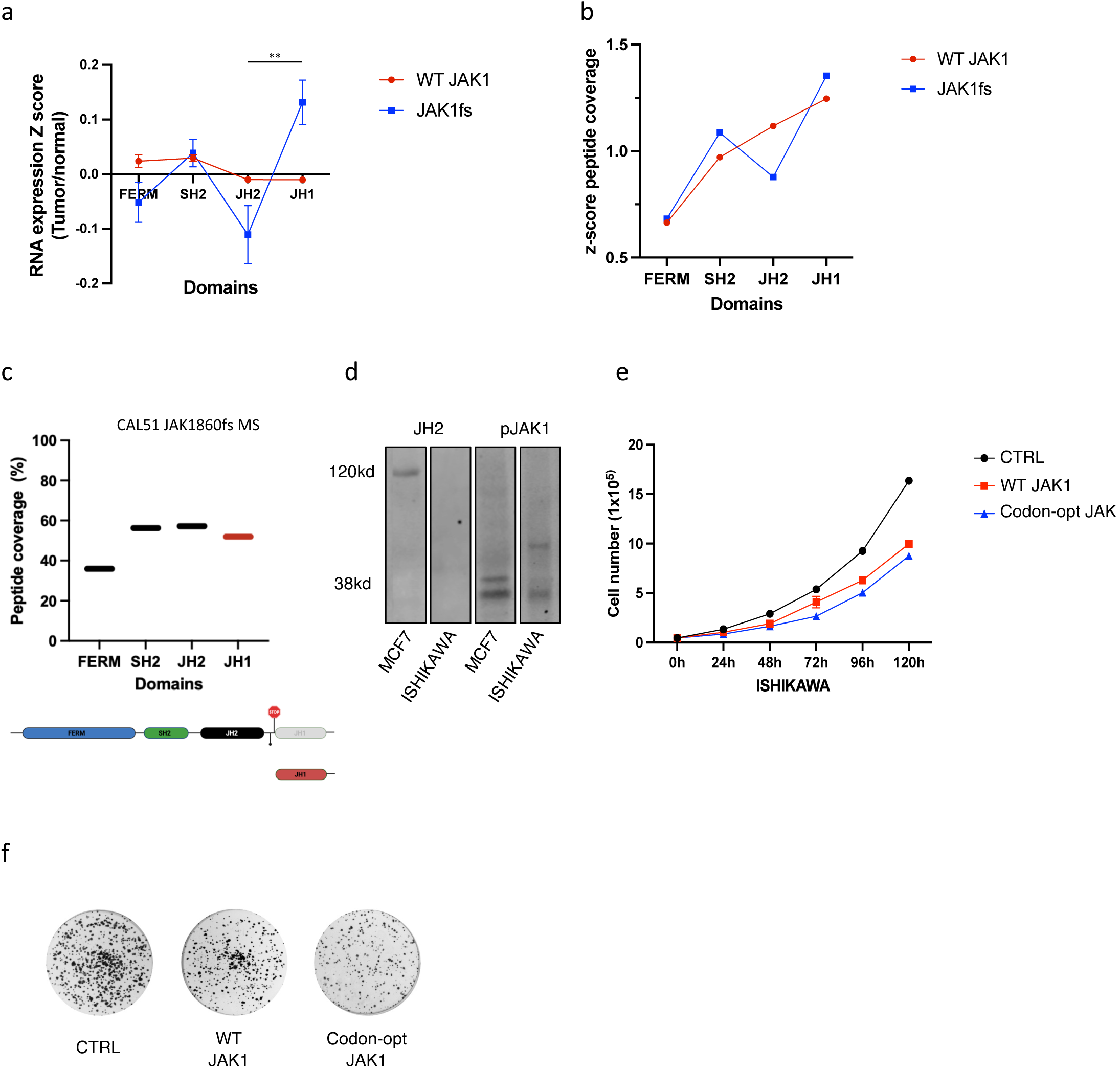
JAK1 frameshift mutations in endometrial cancer affect the expression of the full-length JAK1 isoform but not the expression of the diced JAK1 kinase domain. (a) RNA-seq data from the TCGA-UCEC cohort, comprising 70 patients with WT JAK1 and 7 patients with the JAK1fs mutation, were analyzed. The y-axis represents the relative change in exon expression levels corresponding to different domains. This was calculated by z-transforming JAK1-specific exonic RPKM values for each sample, then subtracting the exon-specific tumor z-scores from the mean z-score of the expression in adjacent healthy tissue for that exon (FERM domain: exons 3-9; SH2 domain: exons 10-12; JH2 domain: exons 13-19; JH1 domain: exons 20-25). The average RNA-seq read coverage deviation between patients and normal adjacent tissue (24 samples) showed no difference in RNA distribution in WT JAK1 patients. However, a sharp decrease in JH2 expression, followed by an increase in JH1 domain expression, was observed in JAK1fs patients, indicating independent JH1 RNA expression. \*\**p* < 0.01 (two-tailed Student’s *t*-test). (b) Mass spectrometry analysis of EC (from the CPTAC cohort) in JAK1fs patients shows a similar trend in domain peptide coverage compared to RNA-seq domain coverage, indicating an independent JAK1 kinase domain isoform in these patients. (c) Mass spectrometry analysis of CAL51 cell line (Cancer Cell Line Encyclopedia) harboring homozygous JAK1fs mutation at position 860 shows 50% in-frame peptide coverage downstream of the frameshift position. (d) Western blot of MCF7 and ISHIKAWA cell lines shows diced phosphorylated JAK1 expression in WT expressing JAK1 MCF7 and ISHIKAWA cells harboring homozygous frameshift mutation at position 860. Full-length JAK1 is only detected in MCF7 cells. (e) Cell growth of cells expressing WT JAK1 or codon-optimized JAK1 constructs was measured using cell counting. Full-length JAK1 expression significantly reduced cell growth. (f) Colony formation assay shows a reduction in colonies in both JAK1 full-length expressing construct cells.

**Figure 5.**
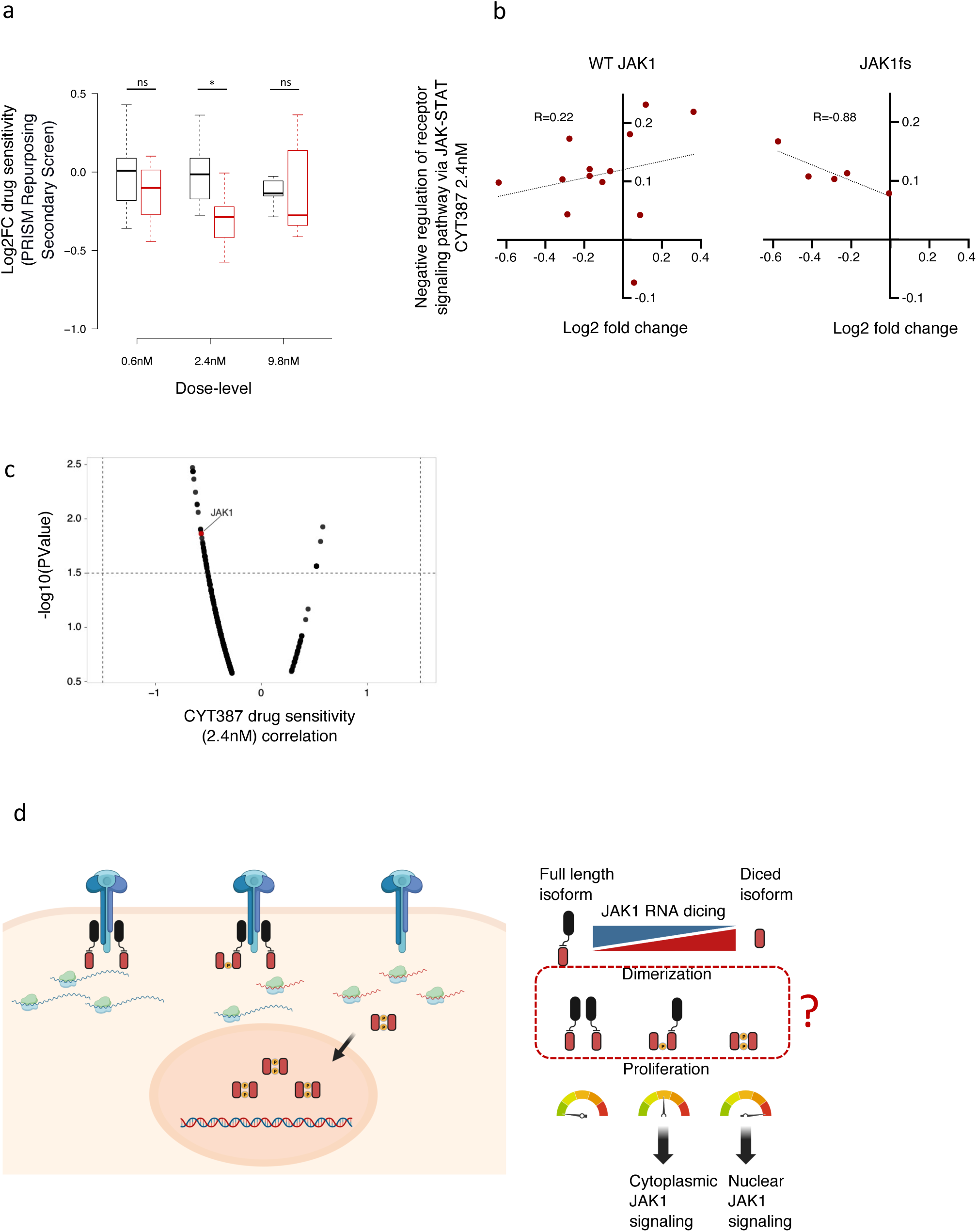
Cells with JAK1fs mutations exhibit high intolerance to JAK inhibitors due to the expression of the oncogenic diced JAK1 kinase and complete KD of the canonical full length tumor suppressor. (a) Drug sensitivity analysis using the PRISM Repurposing Secondary Screen of the JAK inhibitor CYT387 at different concentrations in multiple EC cell lines, including 19 cell lines expressing wild-type JAK1 (black) and 5 cell lines with homozygous JAK1fs mutations (red) \**p* < 0.05 (two-tailed Student’s *t*-test). (b) Single-sample gene set enrichment analysis was conducted for the GO category ‘Negative Regulation of Receptor Signaling Pathway via JAK-STAT’ following treatment with CYT387 at a concentration of 2.4 nM in WT and frameshift JAK1 EC cell lines. (c) Correlation between damaging mutation and CYT387 drug sensitivity. Out of 17 EC cell lines that were treated with CYT387 (2.4 nM), cells harboring a JAK1 mutation were found to be among the most sensitive to this treatment. This included 9 cell lines with WT JAK1, 3 cell lines with a heterozygous mutation, and 5 cell lines with a heterozygous damaging mutation in JAK1. (d) **JAK1 Dicing Model:** The JAK1 protein can be expressed as both the full-length canonical isoform and a diced isoform, which primarily includes the JH1 kinase domain. JAK1 activation is facilitated by dimerization and trans-phosphorylation through the JH1 kinase domain. We postulate that this enables the potential formation of both heterodimers and homodimers. Increased JAK1 dicing shifts the stoichiometry from the canonical to the diced JAK1 isoform, thereby increasing the likelihood of forming canonical-diced or diced-diced dimers. The absence of the FERM and JH2 domains removes the isoform’s restriction to the cell membrane and enhances phosphorylation and activation due to the loss of the JH2 pseudokinase domain. This shift enables alternative signaling pathways downstream of JAK1, which are physically located in the nucleus which accelerates cell cycle.

Finally, we extended this analysis to two other cancer cells with known homozygous JAK1860fs mutations (CCLE). The Invasive Breast Carcinoma cell line CAL51 was more sensitive to CYT387 than any of the 19 other cell lines from the same tumor type that were tested. Interestingly however, the IGROV1 EC cell line did not show any response to CYT387 treatment (Extended Data Fig.5d). However, MS analysis suggests that while the CAL51 cell line expresses the JH1 domain (Fig.4c), IGROV1 does not, explaining its lack of sensitivity to CYT387 (CCLE, in comparison to 9 other cell lines, Table 2). These findings indicate differential regulation of dicing/translation of JH1 kinase domain in JAK1fs mutation cancer cell lines. Additionally, these findings underscore the robust efficacy of the JAK1 inhibitor CYT387, in treating different types of cancer with JAK1 nonsense mutations that exhibit expression of the JH1 diced isoform, offering a promising avenue for tailored cancer treatment strategies.

## DISCUSSION

In this paper we introduce a mechanism termed ‘RNA dicing’, and show that in the JAK1 mRNA, dicing generates an uncapped sub-template for protein synthesis. The translation of this diced mRNA results in a JH1 kinase domain, independent of other domains. Our findings underscore a key role of RNA dicing in modulating the phosphorylation dynamics, localization, and function of JAK1.

RNA dicing diversifies protein isoform production, and offers profound insights into gene function and potential. The regulation of dicing appears to be crucial for the varied biological functions mediated by JAK1. Our previous studies have demonstrated that the generation of 5’ UTR variants through dicing, a phenomenon observed across thousands of genes, is intricately regulated by mechanisms such as alternative polyadenylation, m6A modification, and RNA structure. These processes are highly dynamic across different biological conditions^40–42^, suggesting that this mechanism of action may extend far beyond JAK1.

Mechanistically, the dicing of JAK1 provides multiple biological advantages. It adapts the kinetic potential of the JH1 kinase domain, by eliminating domains that restrict kinase activity such as the upstream JH2 pseudokinase domain. The removal of the FERM domain alters cellular localization, preventing membrane targeting. Furthermore, the reduced molecular mass of the diced isoform, which is less than 40kD, allows passive nuclear diffusion^43^, enabling the activation of alternative signaling pathways.

Previous studies have demonstrated that JAK kinases are unlike other tyrosine kinases, which require activation through phosphorylation by an upstream kinase. Instead, independent JAK kinase domains possess intrinsic catalytic activity^15,44,45^. Crucially, that catalytic activity is up to 100-fold higher when the kinase domain is expressed independently, compared to the canonical full-length JAK1^15,44^. Our study reveals that this hyperactive kinase activity can be endogenously induced through the dicing of JAK1 and the independent expression of the JAK1 kinase domain.

The current model of JAK1 activation suggests that transphosphorylation and subsequent activation of JAK1 are facilitated by the structural flexibility of the kinase domain^46^, which occurs in response to cytokine signaling, however this model is largely speculative. Our findings suggest that inhibition of JAK1 dicing leads to increased full-length JAK1 expression, which in turn results in reduced phosphorylation following IFNγ induction (Fig.2c). This observation challenges the conventional view of activation that is dependent on homodimerization of the canonical form of JAK1. The pronounced dicing observed upon macrophage activation (a process highly dependent on JAK1) underscores its significance. These findings may indicate a mechanism in which heterodimerization of JAK1 isoforms may play a role, potentially circumventing the structural constraints imposed by the pseudokinase-pseudokinase interactions of full-length homodimers. Potentially, the various dimer combinations may induce alternative signaling outcomes that could activate either canonical or non-canonical pathways, as well as inhibit and terminate these signaling cascades (Fig.5d).

In the context of cancer, this study also offers crucial insights, particularly into genomic alterations. We show that what is traditionally considered a loss-of-function mutation (a nonsense mutation in JAK1), can have selective effects on different isoforms. Indeed, we show that rather than being less sensitive to a JAK1 inhibitor, cells carrying this mutation are actually more sensitive, due to altered isoform abundance caused by the mutation. High-resolution analysis of dicing-dependent isoform expression may play a significant role in patient stratification and personalized medicine as a result. Specifically, we may need to revisit the classification of nonsense mutations, as these may act as diced-dependent gain-of-function mutations. Such understanding is vital for deciphering the potential complex role of dicing in cancer biology, as these observations may well extend beyond JAK1 mutation.

Previously, we spotlighted the role of RNA metabolism across thousands of genes, contributing to a diverse uncapped RNA fragments derive from the 3’UTRs and CDS, increasing the immunopeptidome repertoire and influencing BCL2 translation regulation. In this study, focusing on JAK1, we show an additional layer of these metabolic processes as RNA dicing, facilitating modular gene function through varied domain expressions. We believe this new paradigm illuminates aspects of gene regulation and functional understanding, paving the way for innovative pharmaceutical approaches based on RNA dicing and modular gene expression.

## Data availability

All sequencing data are available through the Gene Expression Omnibus^3^ under the following accession: icSHAPE GSE117840; HL-60 Long-read RNAseq GSE219923; m6A RIPseq GSE112795; all other sequencing data GSE149204.

## Acknowledgments

YM and FA are supported by the KWF Kankerbestrijding (13878).

## Contributions

Y.M. conceived the project, designed and performed experiments, analyzed data, and wrote the manuscript; W.J.F. contribute funding and wrote the manuscript. R.vd.K. performed experiments; S.J. and C.L. performed mass spectrometry analysis. F.A. analyzed data.

## Methods

### Cell culture

HeLa, MCF7, ISHIKAWA and HL-60 cell lines were grown in Dulbecco’s modified Eagle’s medium, supplemented with 10% (HL-60 with 20%) fetal calf serum, 100 units/ml penicillin, and 100 μg/mL streptomycin at 37°C. Cell lines were regularly tested for Mycoplasma contamination. Cell lines were authenticated by expression analysis based on RNA-seq.

### Terminator phosphate-dependent TEX treatment

For RNA-seq analysis, DNase I-treated and poly(A)-selected RNA was subjected to treatment with Terminator 5ʹ-Phosphate-Dependent Exonuclease (Epicentre; TER51020) following the manufacturer’s instructions. The reaction was then deactivated, and RNA was subsequently purified using the RNA Clean-Up and Concentration MICRO-Elute Kit (Norgen; 61000).

### Colony Formation

In a 6-well plate, 3250 cells were seeded per well. Media was refreshed after 4 to 5 days, and the experiment was concluded on day 10/11 after seeding. The cells were washed with PBS and fixed using 3.7% formaldehyde (Sigma). Subsequently, the cells were stained with 0.1% Crystal Violet. All experiments were performed with three biological replicates.

### Western blotting

Cell lysates, protein quantification and SDS-PAGE were performed as previously described^1^. Fractionations were run on 4-15% gradient gels (TGX, Bio-Rad) using the Laemmli buffer system and blotted on nitrocellulose (pore size 0.2 µm; Pall). Blots were routinely blocked with 5% nonfat dried milk diluted in TBST for 1 h and then, depending on the manufacturer, incubated with primary antibodies for 1 h or overnight.

Subsequent staining was performed with the appropriate LI-COR secondary antibodies. Visualization was performed by use of an Odyssey infrared scanning device (LI-COR). All experiments were performed with either two or three biological replicates. Primary antibodies: HSP 90α/β (Santa Cruz Animal Health sc-13119, 1:1000), JAK1 JH2 domain (Merck 05-1154, 1:500), Phospho-JAK1 (Cell signaling 3331, 1:1000), Phospho-JAK1 (Cell signaling 74129, 1:1000), Non-Phospho JH1 JAK1 (Abcam ab47435, 1:1000). Secondary antibodies: IRDye® 800CW Goat anti-Rabbit IgG (LI-COR 926-32211, 1: 10000) and IRDye® 680RD Goat anti-Mouse IgG (LI-COR, 926-68070, 1:10000).

### Nuclear/Cytoplasmic RNA Extraction

For nuclear and cytoplasmic protein extraction, two million MCF7 cells were utilized. The NE-PER Nuclear and Cytoplasmic Extraction Kit (Thermo Fisher Scientific, Catalogue Number 78833) was employed for this procedure.

### Lentiviral Production and Transduction

To produce lentivirus, 4 × 10^6 HEK293T cells were seeded per 100-mm dish one day prior to transfection. For each transfection, 10 μg of the pCDH reporter, 5 μg of pMDL RRE, 3.5 μg pVSV-G, and 2.5 μg of pRSV-REV plasmids were mixed in 500 μL of serum-free DMEM. Next, 500 μL of serum-free DMEM containing 63 μL of a 1 mg/mL PEI solution was added. The entire mix was vortexed and left for 15 minutes at room temperature, after which it was added to the HEK293T cells for transfection. The following day, the medium was replaced with RPMI. Lentivirus-containing supernatants were collected 48 and 72 hours after transfection and snap-frozen in liquid nitrogen. Target cells were transduced on two consecutive days by supplementing the lentiviral supernatant with 8 μg/mL polybrene (Sigma). One day after the final transduction, transduced cells were selected by adding 2 μg/mL puromycin to the medium.

### Real-time PCR

For reverse transcription, 1 μg of total RNA was utilized and reverse transcribed using the Tetro cDNA synthesis kit in accordance with the manufacturer’s instructions. Subsequently, real-time PCR was conducted using the SensiFAST SYBR real-time PCR kit (Bioline). The obtained data were normalized to the human endogenous control (GAPDH) and analyzed using the ΔΔCt model, unless stated otherwise. All experiments were performed with either two or three biological replicates.

GAPDH:

F: ACAACTTTGGTATCGTGGAAGG.

R: GCCATCACGCCACAGTTTC

JAK1:

UP:

F: TGACGAGAACACCAAGCTCT

R: GAGAATGACGCCACACTGAC

DOWN:

F: TGCACAGAAGACGGAGGAAA

R: GAACGTATTGCCGAGAACCC

JAK1 (Constructs specific to WT and APA-mut JAK1):

UP:

F: CAAAAAAGCAGGCTGCCAC

R: GTGTACTCTCCACTGCCCAG

DOWN:

F: GCCCACCTAACTGTCCAGAT

R: CTTTGTACAAGAAAGCTGGGTT

JAK1 (Constructs specific to Codon optimized JAK1):

UP:

F: TCCGAGACCCTAAAACCGAG

R: CAGGTTCCGTTGTCTGATGC

DOWN:

F: TGGGAGATCTGCTACAACGG

R: GCATAATGGCCCGGAAGAAG

### siRNA knock down

siRNAs against JAK1 (UP and DOWN) were purchased from Life Technologies catalog number AM16708 and 4390824, ID 242351 and s7648. Negative control siRNA catalog number 4390844. Cells were transfected for using Dharmafect I reagent (Dharmacon) following the manufacturer’s instructions (2 siRNAs per target gene) for 72h.

### 5ʹ End Sequencing

20 μg of total RNA was polyA-selected and then treated with E. coli-purified AlkB for demethylation of the RNA, as previously described^2^. Next, we constructed uncapped 5ʹ-specific sequencing libraries, as previously described^3^. The experiment was performed with two biological replicates.

### Illumina RNA-sequencing

RNA was extracted from HeLa cells using QIAzol Reagent (15,596-018, Ambion life technologies) according to the manufactures protocol followed by DNase I treatment. PolyA selected RNA was isolated using Oligotex kit (QIAGEN) and further processed with SMARTer Stranded RNA-Seq Kit (Takara; 634,839) and illumina Truseq Stranded mRNA Library Prep kit. or 3ʹ-mRNA-Seq Library Prep Kit (lexogen; 015UG009V0211).

### RNA-Seq analysis

#### Trimming and filtering of raw reads

NextSeq basecall files were converted to FASTQ files using the bcl2fastq (v.2.15.0.4) program with default parameters.

#### QC preprocessing

Raw reads were inspected for quality issues with FastQC (v.0.11.2) and were quality trimmed at both ends to a quality threshold of 32. Adapter sequences were then removed using cutadapt (version 1.7.1) through the Trim Galore! interface (version 0.3.7), leaving only reads of length above 15 nt. The remaining reads were further filtered to remove very low-quality reads, using the fastq_quality_filter (FASTX package, version 0.0.14), with a quality threshold of 20 at 90% or more of the read positions.

#### Genomic mapping of RNA-seq data

The processed FASTQ files were mapped (using TopHat, v.2.0.13)22 to the human genome and transcriptome (hg19). Reads that, after processing, were left as a pair, as well as reads for which only one of the pair mates remained, were used for further analyses. Mapping allowed up to 2 mismatches per read, a maximum gap of 5 bases, and a total edit distance of 7.

#### Nanopore RNA Sequencing

Poly(A) selected RNA (500 ng) from control or TEX treated samples was prepared for nanopore direct RNA sequencing, generally following the ONT SQK-RNA002 kit protocol, which includes the optional reverse transcription step recommended by ONT. RNA sequencing on the MinION was performed using ONT R9 Flow Cells. The experiment was conducted with two biological replicates.

### 3ʹ-End RNA-Seq Analysis

#### Trimming and Filtering of Raw Reads

NextSeq basecall files were converted to FASTQ files using bcl2fastq (v.2.17.1.14). Reads were screened and preprocessed similarly to the described process above. An additional step involved removing polyA sequences from the 3ʹ ends of reads, which was carried out with cutadapt. A 75-mer oligo-A sequence was used as the “adapter,” and a minimal overlap of 2 was required for the removal process.

### Mass Spectrometry

#### Data Sets

Two publicly available TMT-based proteomic spectrum files were downloaded and used for the peptide quantitation analysis of JAK1. The uterine corpus endometrial carcinoma (UCEC) dataset with the CPTAC study identifier of PDC000125^4^, and The Cancer Cell Line Encyclopedia (CCLE) dataset with MassIVE identifier of MSV000085836^5^.

#### Databases

The concatenated protein database of human reference proteome database from UniProt (Release 2023_01, 20,603 entries) and universal contaminant database^6^. To false discovery rate control purpose, the protein database was attached decoy sequences via FragPipe proteome search program (v19.1).

#### Protein Identification and Quantification

The built-in workflow “TMT10” was used with the adjustments described next. For CCLE dataset, the downloaded raw files were converted to mzML using the ProteoWizard MSConvert tool (v3.0.20287) and for UCEC dataset, we used the mzML files without conversion. For MSFragger (v3.7)^7^ settings, digestion enzyme, trypsin; number of tolerable termini (Cleavage) was set 1 (SEMI) with clipping N-terminal methionine; variable modification, M: 15.9949 (Oxidation), protein N-term.: 42.0106 (Acetyl), peptide N-term. and S: 229.16293 (TMT); fixed modification, C:57.02146 (Carbamidomethyl), K:229.16293 (TMT). For validation settings, MSbooster (v1.1.11) was used for RT and spectra prediction; PSM validation: Percolator (v3.05)^7^ with tdc option with minimum probability of 0.5. For isobaric quantification, TMT-10 label type was selected and reference channel was set as TMT-126 channel for both UCEC and CCLE dataset, normalization was done at peptide level of median centering, minimal accepted purity was 0.75.

#### Data Interpretation

For UCEC, we selected 7 patients with JAK1 mutation of 860 frame-shift: C3N-00850-02, C3L-01744-01, C3N-00389-04, C3N-01212-03, C3L-01257-01, C3N-00321-01, C3N-01219-03. For CCLE, we selected CAL51, SNUC2A, IGROG1, HEC265, HEC108, ISHIKAWAHERAKLIO02ER, 22RV1, MFE319, SNU1 and MFE296. In CCLE dataset, CAL51 has 3 replicates and we used all. The median normalized peptide level ratio acquired via FragPipe were used to generate JAK1 peptide level relative quantitation data.

