## Supplementary Figures for "RNA dicing regulates the expression of an oncogenic JAK1 isoform"

### Supplementary Figure 1

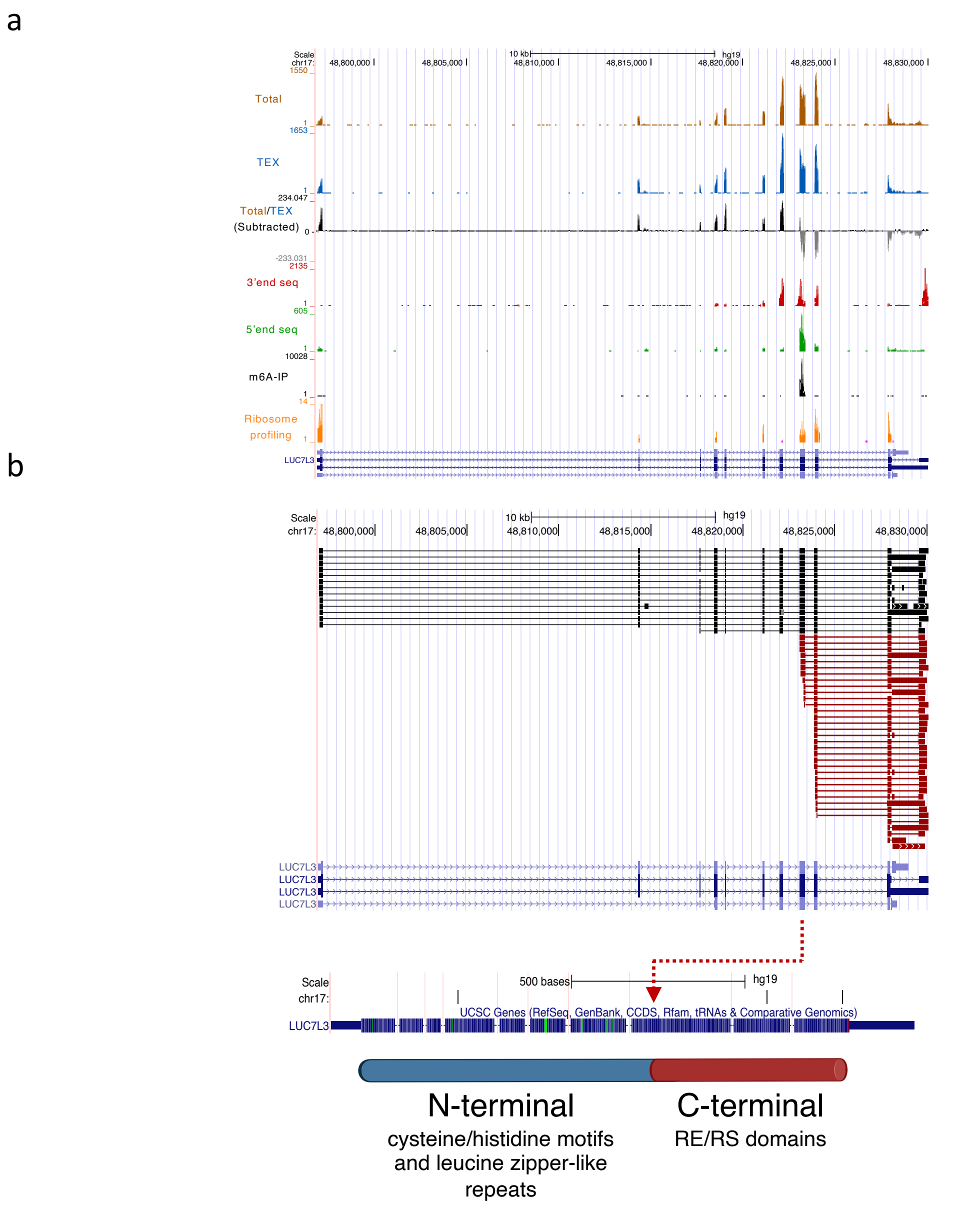


```
MISAAQLLDELMGRDRNLAPDEKRSNVRWDHESVCKYYLCGFCPAELFTNTRSDLGP
CEKIHDENLRKQYEKSSRFMKVGYERDFLRYLQSLLAEVERRIRRGHARLALSQNQQ
SSGAAGPTGKNEEKIQVLTDKIDVLLQQIEELGSEGKVEEAQGMMKLVEQLKEEREL
LRSTTSTIESFAAQEKQMEVCEVCGAFLIVGDAQSRVDDHLMGKQHMGYAKIKATVE
ELKEKLRKRTEEPDRDERLKKEKQEREEREKERERERERERKRRREEEEREKERAR
DRERRKRSRSRSRHSSRTSDRRCSRSRDHKRSRSRERRRSRSRDRRRSRSHDRSERK
HRSRSRDRRRSKSRDRKSYKHRSKSRDREQDRKSKEKEKRGSDDKKSSVKSGSREKQ
SEDTNTESKESDTKNEVNGTSEDIKSEGDTQSN
```

### Supplementary Figure 1

a) Multi-omics analysis of LUC7L3 overview displays concentration of endonuclease cleavage, resulting in termination/generation of uncapped truncated RNA. RNA-seq read coverage (y-axis) across the putative cleavage point of LUC7L3 shows reduced downstream read coverage in TEX (exonuclease) and 5prime (capped RNA pulldown) treated cells compared to enriched downstream read coverage in 3prime (uncapped RNA pulldown), as compared to coverage in untreated RNA (Total). APA (3'end seq) induces endonuclease cleavage, 5'end seq indicates 5'end of uncapped RNA and m6A-IP m6A site. Ribosome profiling after harringtonine treatment indicate ribosome translation initiation sites

b) Nanopore long RNA sequencing data, displayed in a genome browser view, show isoforms aligned to LUC7L3. Black isoforms are adjacent to the promoter region at their 5' ends, and red isoforms represent 5' end diced RNA corresponds to the C terminal coding region of RE/RS domain.

### Supplementary Figure 2

a

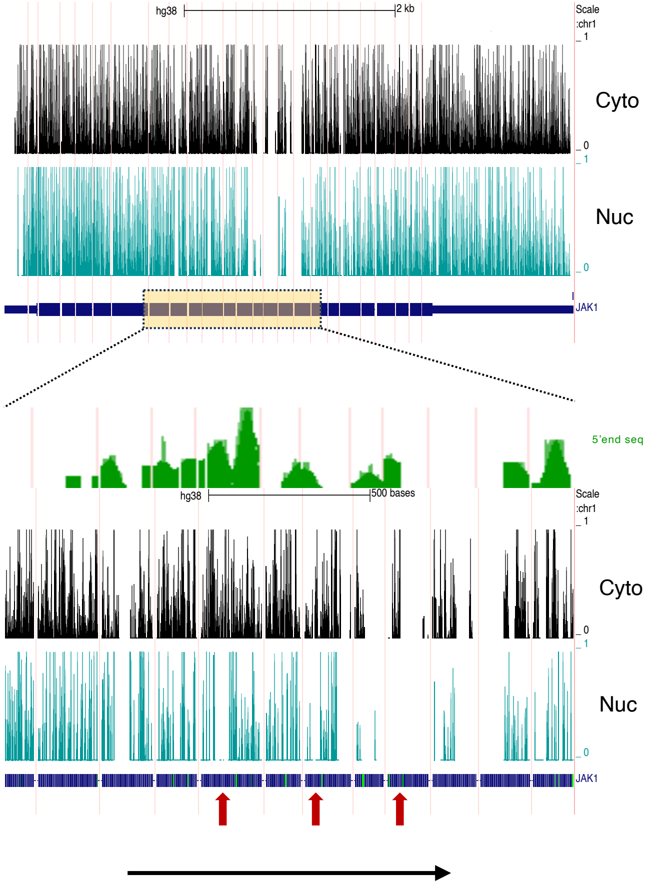


b

JAK1 Exon 8 (chr1:65330470-65330655 hg19)
GTTGTTTCTGTTGAAAAGGAAAAA**AATAAA**CTGAAGCGGAAAAAACTGGA
A**AATAAA**CACAAGAAGGATGAGGAGAAAAACAAGATCCGGGAAGAGTGGA
ACAATTTTTCTTACTTCCCTGAAATCACTCACATTGT**AATAAA**GGAGTCT
GTGGTCAGCATTAACAAGCAGGACAACAAGAAAATG

c

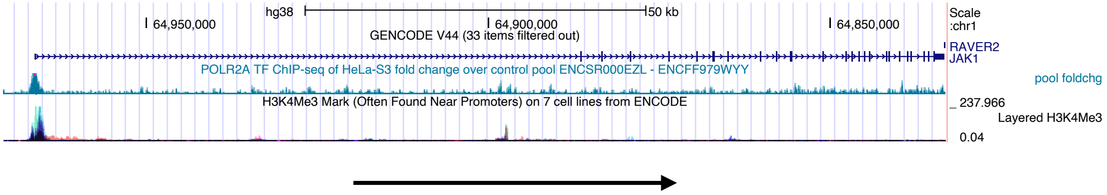


d

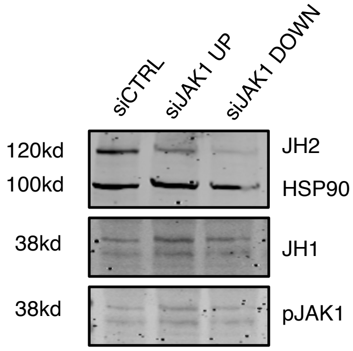


e

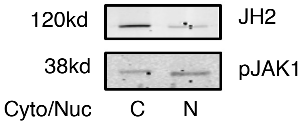


f

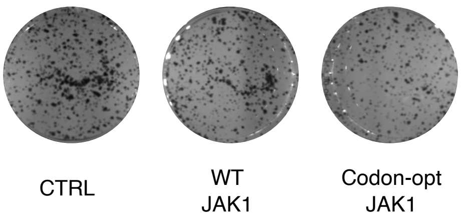

### Supplementary Figure 2

a) Analysis of icSHAPE data (in vivo and in vitro, HEK293 cells) presents a strong prediction of double-stranded RNA around the 5'end of diced uncapped RNA. The magnified lower panel shows predicted structured RNA corresponding to 5'seq. Structured RNA in the cytoplasmic fraction compared to the nuclear fraction (red arrows) indicates post-transcriptional dicing.
b) Exon 8 of JAK1 contains several conserved APA signal sites (AATAAA).
c) GROseq analysis of JAK1 indicate single transcription initiation at the canonical site correspond the H3K4Me3 histone site.
d) Western blot analysis of MCF7 cells following siRNA transfection against JH2 JAK1 (UP) or JH1 domain (DOWN).
e) Western blot analysis of total JAK1 or phospho-JAK1 from MCF7 cytoplasmic/nuclear fraction lysate.
f) Colony formation by JAK1-expressing constructs (biological replicate of Fig.2h).

### Supplementary Figure 3

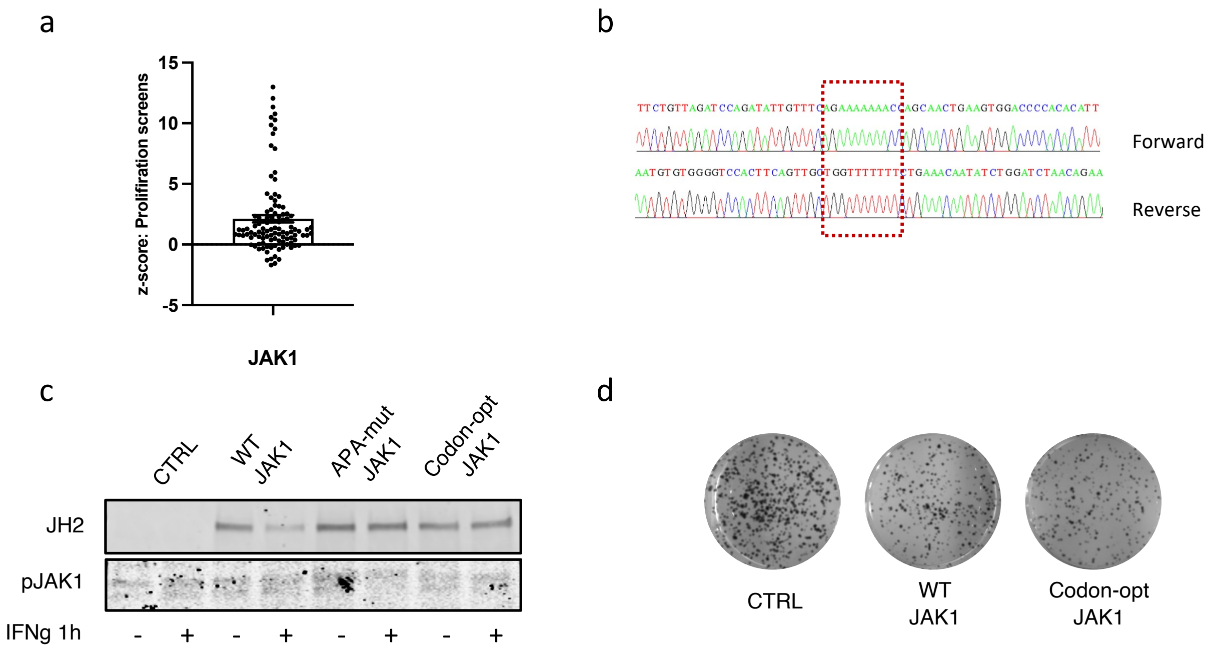

### Supplementary Figure 3

(**a**) Z-scores from 108 base editing proliferation screens inducing nonsense stop codon mutations are plotted for each gRNA. Screen z-scores are calculated independently for each base editor and plotted together for comparison.

(**b**) Sanger sequencing of JAK1 in the ISHIKAWA cell line, focusing on the region surrounding the frameshift site at position 860, revealed seven adenosines instead of the eight found in the wild type.

(c) Western blot analysis was performed on ISHIKAWA cells expressing different JAK1 constructs.

(d) Colony formation by JAK1-expressing constructs (biological replicate of Fig.4f).

### Supplementary Figure 4

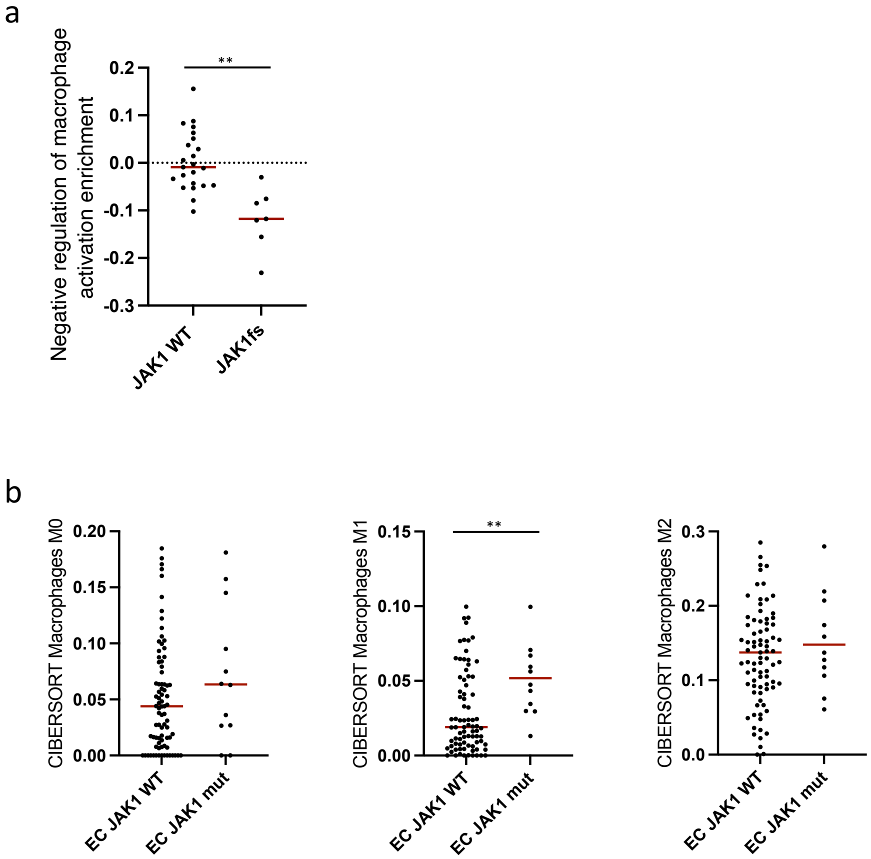

Supplementary Figure 4

(**a**) Single-sample gene set enrichment analysis of EC cell lines with different JAK1 mutation status shows decrease in negative regulation of macrophage activation enrichment in JAK1fs mutated cells.

(**b**) Single-sample gene set enrichment analysis of EC patients (CPTAC cohort) shows an increase in the M1 macrophage signature in patients with a JAK1fs mutation status. $^{**}p < 0.01$ (two-tailed Student's *t*-test).

### Supplementary Figure 5

a

Response of multiple cell lines to CYT387 at different concentrations in relation to JAK1 RNA levels.

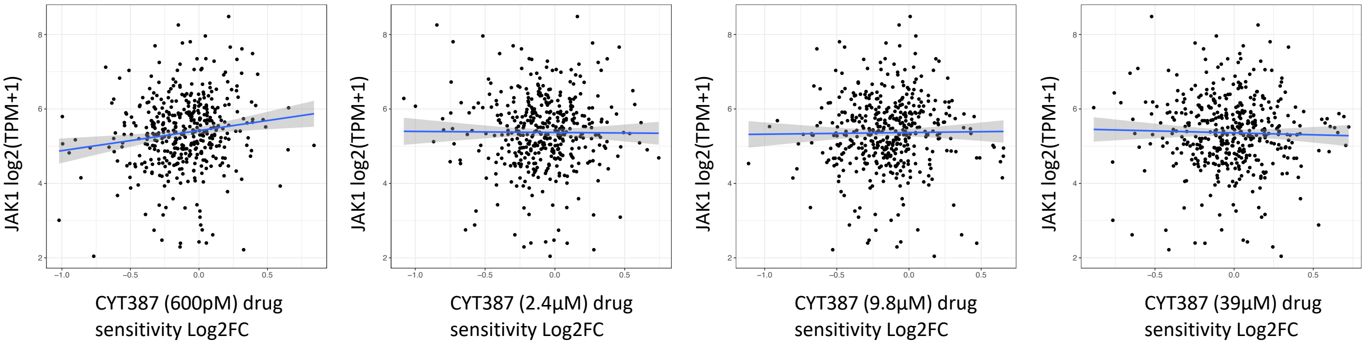


b

Response of multiple cell lines to CYT387 at different concentrations in relation to JAK1 protein levels.

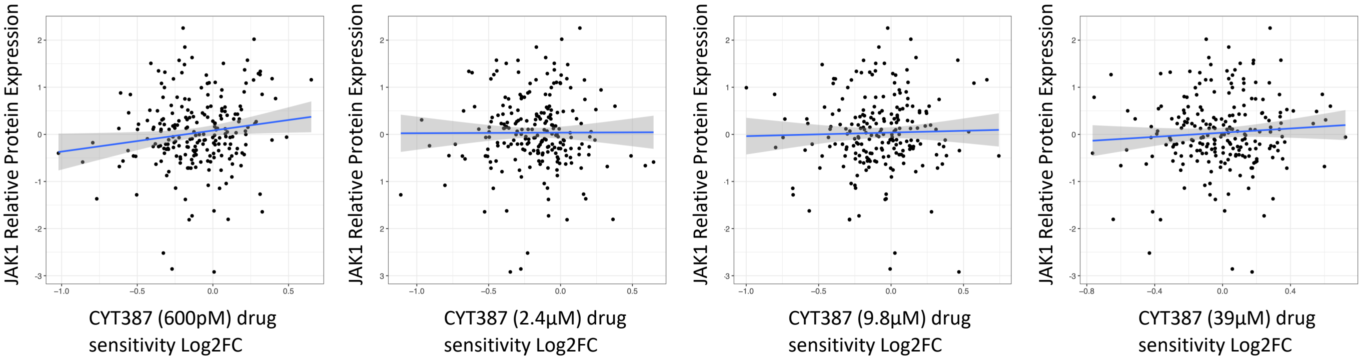


c

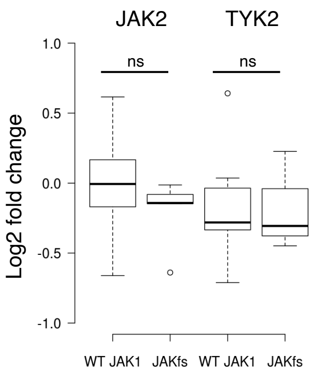


d

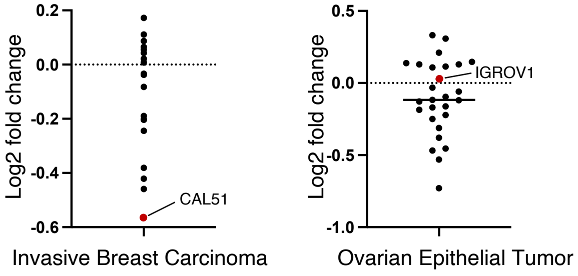

### Supplementary Figure 5

(**a-b**) RNA and protein expression of JAK1 (434 and 221 cell lines respectively) showed no correlation with the efficacy of CYT387 treatment at various doses.
(**b**) Protein level of JAK2 and TYK2 in EC cell lines with different JAK1 mutation status (JAK2 - protein array analysis; TYK2 – proteomic analysis).
(**c**) Cell lines from invasive breast carcinoma and ovarian epithelial tumor, carrying either homozygous JAK1860fs mutations or WT JAK1, and their response to CYT387 (2.4nM).
